# Preservation Strategies for Vascularized Composite Allotransplantation: An Updated Systematic Review of a Rapidly Expanding Field

**DOI:** 10.1101/2025.09.11.675673

**Authors:** Pharel Njessi, Pierre Barbat, Rabbani S Piul, Didier F Pisani, Olivier Camuzard, Antoine Sicard, Eduardo Rodriguez, Elise Lupon

**Affiliations:** Department of Plastic and Reconstructive Surgery, Institut Universitaire Locomoteur et du Sport, Pasteur 2 Hospital, University Côte d’Azur, Nice, France; Université Côte d’Azur, CNRS, LP2M, France; Faculty of Medicine, University of Montreal, Montreal, QC, Canada; Hansjörg Wyss Department of Plastic Surgery, New York University Langone Health, New York; Department of Nephrology, Dialysis and Kidney Transplantation, University Hospital of Nice, Nice

## Abstract

**Background:** Vascularized composite allotransplantation has become a viable reconstructive option for selected patients, but preservation remains a major barrier to broader clinical application. Static cold storage is the current gold standard, yet ischemia reperfusion injury and limited preservation times restrict its effectiveness. Recent advances in machine perfusion and subzero non-freezing storage (or supercooling) have prompted renewed interest in optimizing graft viability.

**Methods:** Following PRISMA guidelines, we systematically searched PubMed, EMBASE, and Cochrane, covering studies published from June 2022 through August 2025 for studies on ex vivo preservation of vascularized composite allotransplantations. Eligible articles included original studies in English evaluating postharvest, pretransplant preservation strategies. Data extracted were study design, preservation methods, perfusates, and primary outcomes. Risk of bias was assessed using SYRCLE for animal studies and JBI for human/cadaver studies.

**Results:** Seventeen studies met inclusion criteria: one on static cold storage, thirteen on machine perfusion, and three on supercooling. Static cold storage research has declined, with the only recent study investigating sub-normothermic machine perfusion as a recovery adjunct. Machine perfusion studies focused on optimization of perfusion parameters, perfusate composition, and circuit design. Red blood cell-based perfusates remained common, but alternative oxygen carriers such as polymerized hemoglobin-based oxygen carrier-201 and dextran oxygen microcarriers showed promise despite edema-related challenges. Supercooling studies demonstrated feasibility of multi-day preservation in rodent and porcine models. Overall, risk of bias was high or unclear across animal studies, mainly due to selection and performance bias, whereas the single human ex vivo study showed low risk of bias.

**Conclusions:** The field of vascularized composite allograft preservation is expanding rapidly, with machine perfusion and supercooling emerging as the most promising strategies to extend graft viability beyond the limits of static cold storage. However, translation to clinical setting remains limited by small preclinical studies, methodological heterogeneity, and the paucity of functional and immunologic endpoints. Standardized protocols, robust large-animal models, and eventual human feasibility trials are needed to establish clinically applicable preservation strategies.

*Level of evidence: IV*

## Introduction

Vascularized composite allotransplantation (VCA) involves the simultaneous transfer of heterogeneous tissues including skin, nerve, tendon, muscle and bone. Over the past two decades, this field has significantly advanced establishing VCA as a viable alternative to conventional reconstructive techniques for congenital, traumatic and oncologic defects. Nevertheless, its widespread application remains limited by challenges, particularly the difficulty in maintaining graft viability (1).

The need for rapid graft reperfusion after procurement represents a major constraint. Currently, static cold storage (SCS) on ice remains the gold standard (2). By lowering cellular temperature to 4 °C, SCS slows metabolic activity and reduce ATP consumption, reducing the imbalance between energy supply and consumption. The objectives of VCA preservation are multiple: to limit ischemic and ischemia–reperfusion injury (3), improve graft quality, allow perfusion of immunomodulatory agents (4), prolong recipient preparation time in tolerance induction protocols using chimerism, to and overcome geographical constraints by enabling long-distance transplantation (5). Optimizing ischemia times is therefore a priority in transplantation, in order to limit inflammatory responses and their harmful consequences (6,7). In the short term, inadequate preservation can lead to degeneration of nerve fibers, thereby compromising the functional prognosis of the graft (8). While some recommend limiting warm ischemia to 6h and cold ischemia to 12h (7), muscle - the most ischemia-sensitive and often predominant in VCAs - can show injury after only 2 hours of warm ischemia or 8 hours of cold ischemia (7,9).

Our group previously published a scoping review of ex vivo pretreatment strategies for VCA including articles available up to June 15, 2022. That review concluded that while innovative preservation techniques showed promise, further investigation was needed to assess long-term outcomes and establish protocols capable of extended preservation times (10). Since then, the field of VCA preservation has experienced a notable surge in experimental research. An updated review is therefore warranted to integrate these recent findings and reassess the state of the field. This systematic review aims to assess the most recent studies devoted to optimizing VCA preservation.

## Methods

### Search strategy

Following PRISMA (Preferred Reporting Items for Systematic reviews and Meta-Analyses) guidelines (11), we conducted a literature search of PubMed, EMBASE, and Cochrane using terms related to VCA and transplantation (Supplementary Material 1). The electronic databases were searched from 2022 to August 2025 for articles published in English. Additionally, we conducted manual searches of reference lists from identified articles.

### Eligibility criteria

Included studies met the following criteria: (1) original text written in English; (2) full-text articles; (3) articles that studied preservation methods on ex vivo VCA, meaning postharvest and pretransplant; (4) articles published between 2022 and August 2025; and (5) articles focusing on cell physiology and graft viability. Exclusion criteria comprised: (1) non-original articles (i.e. abstracts, reviews, commentaries, etc.); (2) non-English articles; (3) articles not reporting primary data; (4) articles not related to VCA transplantation (i.e. articles on other organs or on flaps without osseous or muscle content).

### Study selection

Two independent reviewers (P.N. and P.B.) screened the titles and abstracts of identified articles to assess their eligibility based on the predetermined inclusion and exclusion criteria. Full-text articles of potentially eligible studies were retrieved and reviewed in detail. Disagreements were resolved by discussion or arbitration by a third reviewer (E.L.).

### Data collection

Data from eligible studies were extracted using a standardized data extraction form. The following information was collected: study characteristics (author, year, and country), study design, preservation strategy characterization (process, solutions, timing, etc.), evaluation metrics, and outcomes. The included studies were then organised in 3 categories: (1) static cold storage; (2) ex vivo machine perfusion (MP); or (3) supercooling.

### Data synthesis and analysis

Given the heterogeneity in study design and outcome measures, a meta-analysis was not favored. Instead, a narrative synthesis of the included studies was conducted. The data were organized and summarized in tabular form, highlighting the characteristics of the included studies, key findings, and outcomes of interest. In addition to extracting clinical and technical variables, we assessed the methodological quality of each included study. Risk of bias was evaluated using SYRCLE’s risk of bias tool for animal studies (12) and JBI critical appraisal checklist for quasi-experimental studies (JBI) for human cadaver studies (13).

## Results

### Study selection

The search strategy is summarized in Figure 1. An initial search of the databases yielded 1275 records related to our keywords. After deduplication, 816 unique records were screened and of those, 44 unique articles were assessed for eligibility. Finally, 17 studies met the predefined inclusion and exclusion criteria and were included in the review. No additional records were identified through manual searches of references.

**Figure 1.**
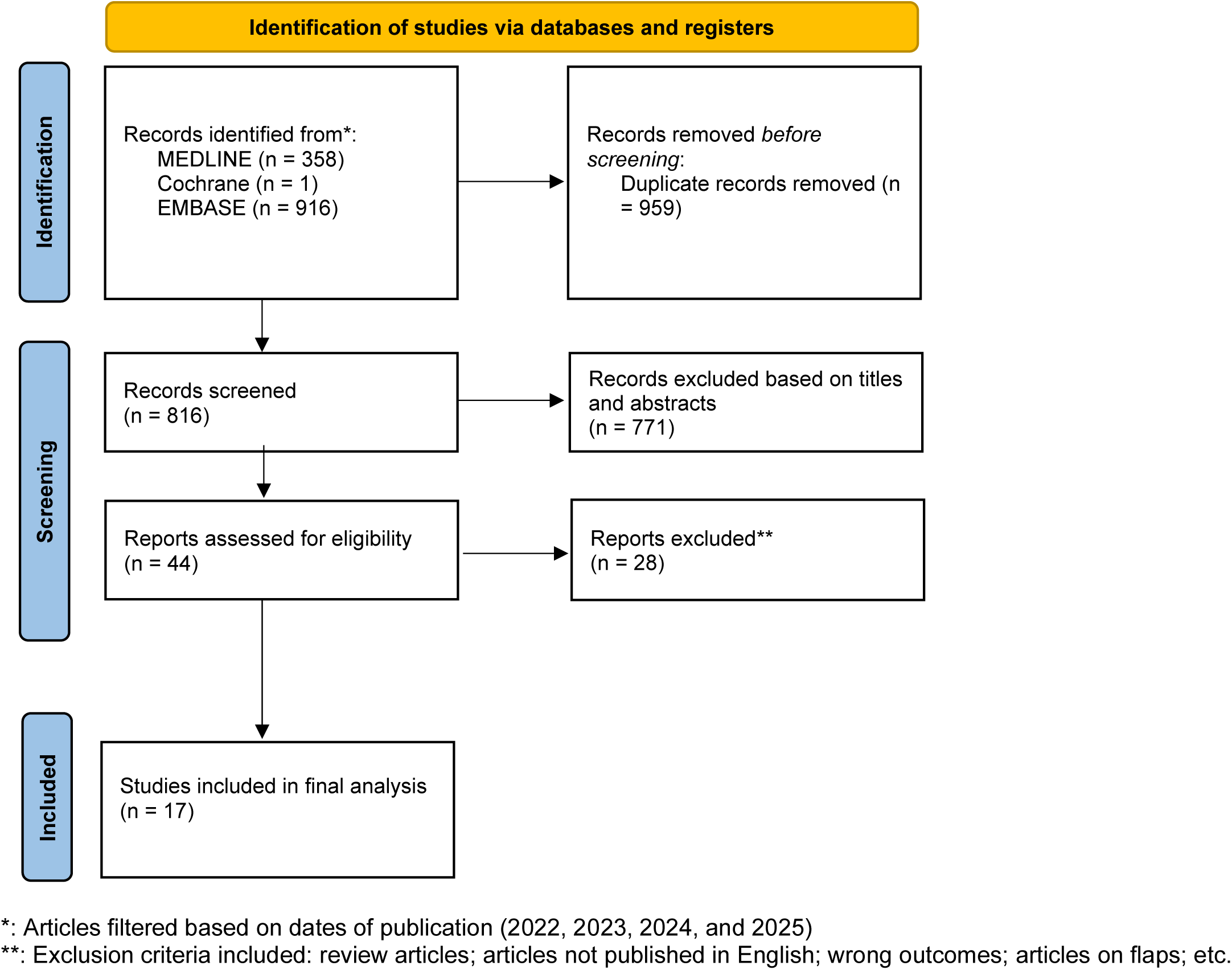
PRISMA Flow chart.

### Study characteristics

Tables 1 and 2 summarize characteristics of included studies. The studies were mainly preclinical and included mostly non-primate animal (rats, pigs and dogs) models with only one cynomolgus monkey model (14) and one human forearm model (15). In many cases, researchers only analysed graft viability without performing transplantation or replantation. Apart from the Brouwers’ study on porcine myocutaneous rectus abdominis flaps (16), all studies used limb models containing skin, blood vessels, muscle, tendon, subcutaneous tissue, fascia, and bones. The preservation methods evaluated included: static cold storage (SCS) (17), machine perfusion (MP) systems (14–16,18–27), and supercooling (28–30).

**Table 1.** Characterizations of Experimental Ex Vivo Perfusion Pretreatment Protocols.

| Study Identifier | Animal Model | Transplant Type | Graft Model | Experimental Groups | Perfusate | No. of Grafts Per Group | Total No. of Grafts | Primary Outcomes |
| --- | --- | --- | --- | --- | --- | --- | --- | --- |
| Brouwers et al., 2022 | Dutch Landrace Pigs | Orthotopic replantation | Rectus abdominis myocutaneous flap | Group 1: flushed with 100 mL heparin-HTK solution + WIT (#1: 38.2°C, 14 min; #2: 37.7°C, 9 min) HMP with modified-HTK (8°C, 24h) + WIT (#1: 14.8°C, 94 min; #2: 12.7°C, 55 min) + Replant<br>Group 2: flushed with 100 mL heparin-UW-MPS + WIT (#1: 37.3°C, 19 min; #2: 38.4°C, 15 min) + HMP with UW (8°C, 24h) + WIT (#1: 13.2°C, 63 min; #2: 13.7°C, 56 min) + Replant<br>Group 3 (control): heparin-saline solution (200E heparin per 100 mL 0.9% saline solution) + WIT (#1: 38.1°C, 5 min; #2: 37.3°C, 5 min) + SCS (4°C, 4h) + WIT (#1: 12.8°C, 62 min; #2: 7.2°C, 34 min) + Replant | Group 1: Oxygenated acellular HTK enriched with PEG and L-glutamine, methylprednisolone, glucose, insulin<br>Group 2: Oxygenated acellular UW, methylprednisolone | 2 | 6 | Machine perfusion: macroscopic outcomes comparable to 4h of SCS<br>HTK: decreased muscle damage on POD 7 after replant compared to UW, significant increase in weight (60% and 97%) after 24h of machine perfusion.<br>UW: worse histological outcomes 7 days after replantation compared to HTK |
| Figueroa et al., 2022 | Yorkshire pig | None | Forelimbs | Group 1: flush with heparinized saline + WIT (35.50 ± 8.62 minutes) + EVNLP with HBOC-201-based perfusate (27-38°C, 22.50 ± 1.71 hours)<br>Group 2: flush with heparinized saline + WIT (30.17 ± 8.03 minutes) + EVNLP with RBC-based perfusate (27-38°C, 28.17 ± 7.34 hours)<br>Group 3 (control): flush with heparinized saline + WIT (37.82 ± 10.45 minutes) + SCS (4°C, 24h) | Base Perfusate: Plasma-Lyte A, insulin, heparin, vancomycin, albumin, calcium gluconate, methylprednisolone +/- dextrose +/- tromethamine. 400-mL exchanges every 3 hours starting at time point 6<br><u>Group 1:</u> Base perfusate + HBOC-201<br><u>Group 2:</u> Base perfusate + washed RBC | Group 1: 6<br>Group 2: 6<br>Group 3: 12 | 24 | HBOC-201-based perfusate: comparable tissue preservation, contractility, and oxygen utilization as RBC-based perfusate<br>Both HBOC-201 and RBC-based perfusates outperformed SCS |
| Rezaei et al., 2022 | Human | None | Forearm (15 cm above the medial epicondyle) | Group 1: flush with UW + NMP (38°C, 41.6 ± 9.4h) in continuous flow<br>Group 2: flush with UW + SCS (4°C, h) | Oxygenated homologous blood, albumin, heparin, vancomycin, cefazolin, methylprednisolone, dextrose, insulin. replacement every 3 h (starting time point 6) | 10 | 20 | Perfused limbs: muscle contraction preserved for a median 30.5 hours; less injury on biopsy; minimal weight gain (0.4% ± 12.2%)<br>SCS limbs: no muscle contraction during entire experiment; |
| Veraza et al., 2022 | Sus scrofa swine | None | Forelimb | Group 1: flush with heparinized saline + SNMP with ULiSSES closed pressurized perfusion device in modified Krebs-Henseleit solution (15–22°C, 24h)<br>Group 2: flush with heparinized saline + SNMP with conventional open perfusion system in modified Krebs-Henseleit solution (15–22°C, 24h) | Oxygenated acellular modified Krebs-Henseleit solution | Group 1: 9<br>Group 2: 4 | 13 | Closed perfusion: 21% reduction in weight gain and significantly reduced intracellular edema; 70% oxygen consumption; lower vascular resistance. Closed pressure → ↓ edema → ↓ vascular resistance → ↓ reducing blood supply → cellular damage and reduced oxygen consumption<br>Open perfusion: marked edema (weight gain +32.9%); 20% oxygen consumption; |
| Tawa et al., 2023 | Yorkshire pig | None | Forelimb | Group 1: flush with heparinized serum + TIT ($15 \pm 4$ min) + SNMP with Steen+ ($21^{\circ}\text{C}$, 24h) in continuous flow<br>Group 2: flush with heparinized serum + TIT ($16 \pm 3$ min) + SNMP with Steen+ ($21^{\circ}\text{C}$, 24h) in pulsatile flow (60 beats/min) | Oxygenated acellular modified Steen+, albumin, PEG, heparin, insulin, dexamethasone, hydrocortisone, penicillin-streptomycin; exchanged after 12h. | 3 | 6 | Pulsatile flow: higher arterial resistances; higher NO/ET-1 ratio, reflecting improved vasoconductivity;<br>Continuous flow: quicker increase in weight; higher lactate levels; no histological difference with pulsatile flow |
| Goutard et al., 2023 | Inbred Lewis rats | Heterotopic transplant | Partial Hindlimbs | First part: VCA Viability after Extended Durations of SCS (0, 12, 18, 24, and 48-Hour SCS+Txp)<br>5 Groups with varying durations: flush with heparinized saline + flush with HTK + SCS in HTK solution ($4^{\circ}\text{C}$) + transplant<br>Second part: Effect of 3-hour SNMP after 0, 12, and 18-hour SCS<br>3 Groups with varying durations: flush with heparinized saline + flush with HTK + SCS in HTK solution ($4^{\circ}\text{C}$) + SNMP with modified Steen+ ($21^{\circ}\text{C}$, 3h)<br>Third part: VCA Transplantation after 12-hour SCS and 3-hour SNMP<br>Group 1: flush with heparinized saline + flush with HTK + SCS in HTK solution ($4^{\circ}\text{C}$, 12h) + transplant<br>Group 2: flush with heparinized saline + flush with HTK + SCS in HTK solution ($4^{\circ}\text{C}$, 12h) + SNMP with modified Steen+ ($21^{\circ}\text{C}$, 3h) + transplant | Oxygenated modified Steen+, albumin, PEG, heparin, insulin, dexamethasone, hydrocortisone, penicillin-streptomycin. | 4 | 60 | First part: Graft survival rate was found similar in the 12, 18, and 24-hour SCS+Txp groups. The survival rate in the 48-hour SCS+Txp group was significantly lower than in other groups<br>Second part: Increased weight gain and ischemic lesions with increasing SCS durations<br>Third part: Better clinical outcomes are found in the 12-hour SCS + SNMP + Txp group at POD 21 ($p = 0.016$) |
| Haug et al., 2023 | Male Sprague-Dawley rats | None | Hindlimb | Group 1: WIT ($22.91 \pm 1.64$ min) + NMP with non-oxygenated dextrane-enriched Phoxilium ($37^{\circ}\text{C}$, 12h)<br>Group 2: WIT ($22.91 \pm 1.64$ min) + NMP with Oxygenated dextrane-enriched Phoxilium ($37^{\circ}\text{C}$, 12h)<br>Group 3: WIT ($22.91 \pm 1.64$ min) + NMP with Phoxilium enriched with dextran oxygen carriers (MO2) ($37^{\circ}\text{C}$, 12h)<br>Group 4: WIT ($22.91 \pm 1.64$ min) + SCS ($+4^{\circ}\text{C}$, 12h) | Perfusate base: modified Steen+ with dextrose, insulin R, methylprednisolone, and heparin +/-<br>Group 1: Non-oxygenated dextrane-enriched Phoxilium<br>Group 2: Oxygenated dextrane-enriched Phoxilium<br>Group 3: Phoxilium enriched with dextran oxygen carriers (MO2) | Group 1: 3<br>Group 2: 3<br>Group 3: 2<br>Group 4: 3 | 11 | Perfusate groups: showed higher tissue $p\text{O}_2$ and lower lactate levels after 8-12h compared to acellular perfusates; weight gain significantly higher in perfusion groups (81-124%) vs. SCS (27%); significant increase in average mitochondrial diameter after 12h<br>SCS: significant decrease in mitochondrial diameter after 12h |
| Ng et al., 2024 | Male Brown Norway rats | None | Hindlimb | Group 1: flush with heparinized KPS-1 + CIT ($4^{\circ}\text{C}$) + HMP with heparinized KPS-1 ($4^{\circ}\text{C}$, 24h) | KPS-1 or NS each containing 4000 units of heparin | 6 | 12 | KPS-1: significantly less edema; significantly less muscle cell death; significantly greater decline in pH |
|  |  |  |  | Group 2: flush with heparinized NS + CIT (4°C) + HMP with heparinized NS (4°C, 24h) |  |  |  | NS: more edema; higher apoptosis; poor structural preservation; similar vascular resistance and flow rate. |
| Chen et al., 2024 | Beagle dogs | None | Forelimb | Group 1: flush with heparinized saline solution + TIT (18 min; 9 to 34 min) + Oxygenated autologous blood (37°C, 24h)<br>Group 2: flush with heparinized saline solution + TIT (18 min; 9 to 34 min) + SCS (4°C, 24h) | Oxygenated autologous blood, hydroxyethyl starch 130/0.4 electrolyte solution, D-calcium gluconate, cefazolin, methylprednisolone, and heparin. Replacement every 6h. | 3 | 6 | NMP: weight gain of 11.7%; lesser muscle damage; increased vascular α-SMA; denser arrangement of muscle fibers.<br>SCS: less weight gain (0.18%); more distortions on muscle biopsies |
| Brouwers et al., 2024 | Female Dutch Landrace pigs | Heterotopic replantation | Forelimb | Group 1: flush heparinized HTK + WIT (34.6°C, 12.9 min) + SNMP with HTK (8°C, 24h) + IT (12.0°C, 58 min) + replantation<br>Group 2: flush heparinized HTK + WIT (35.4°C, 2.6 min) + SCS (4°C, 4h) + IT (6.5°C, 52 min) + replantation | Oxygenated acellular HTK solution, methylprednisolone, glucose, insulin, PEG, and L-glutamine. | 8 | 16 | SNMP: weight gain of 44%; more edema; higher HISS of 3.8<br>SCS: weight decrease of 2%; less edema; HISS of 1.8 |
| Zhang et al., 2024 | White pigs | Orthotopic replantation | Forelimb | Single Group: WIT (22°C, 2h) + flush with hydroxyethyl starch solution + WIT (22°C, 7h) + HMP with modified HTK solution (8-10°C, 24h) + replantation | Oxygenated modified HTK, heparin, L-glutamine, glucose, methylprednisolone, insulin, PEG, autologous washed red blood cells, sodium bicarbonate, amoxicillin. | 6 | 6 | All limbs successfully replanted after 33 h total ischemia. Stable systemic markers. Edema on histology but no myofiber necrosis, preserved muscle integrity. |
| Charlès et al., 2024 | Male Lewis Inbred Rats | Syngeneic partial hindlimb heterotopic transplant | Partial Hindlimb | **Step 1:** 6 Groups with varying WIT (30, 45, 60, 120, 150, and 210 min) in water bath at 28°C followed by 3h of SNMP<br>All Groups: flush with heparinized saline + WIT (28°C)<br>**Step 2:**<br>Group 1: flush with heparinized saline + WIT in saline bag in water bath (28°C, 120 min) + SNMP (23±2°C, 3h) + Txp<br>Group 2: flush with heparinized saline + WIT in saline bag in water bath (28°C, 120 min) + Txp | Oxygenated acellular modified Steen+, albumin, PEG, dextran, heparin, insulin, dexamethasone, hydrocortisone, and penicillin-streptomycin. No recirculation. | Step 1: 3<br>Step 2: 5 | 28 | SNMP prolongs WIT to a maximum of 120 min<br>WIT+SNMP+Txp: significantly better clinical and histological outcomes, even 21 d after surgery.<br>WIT+Txp: 80% graft skin necrosis, myocyte death, endothelial permeability, and cellular infiltration |
| Goutard et al., 2024 | Female Yorkshire Outbred Pigs | Heterotopic allogeneic transplantation | Partial hindlimb | Group 1: flush with heparinized saline + WIT (14.7 +/- 1.5 min) + SNMP with Steen+ (+21 °C, 24h) + WIT (105 +/- 5.5 min) + Transplant<br>Group 2: flush with heparinized saline + WIT (12.3 +/- 4.1 min) + flush with HTK + SCS (+4°C, 24h) + flush with saline + WIT (94.7 +/- 10.4 min) + Transplant | Oxygenated acellular Steen+, albumin, PEG, heparin, insulin, dexamethasone, hydrocortisone, and penicillin-streptomycin. 1L exchanged after 12h | 3 | 6 | SNMP: 3/3 in good shape throughout the study.<br>SCS: 2/3 showed deterioration in general condition → euthanized at POD 5 and 9; worse histological findings (variations in myocytes shape and size, and damaged muscle fibers); massive edema and seroma formation |
| Oubari et al., 2025 | Nonhuman primate (cynomolgus) | None | Left forearm | Group 1: flush with heparinized saline + WIT (180 ± 10.95 min) + SNMP with Steen (+19-21°C, 24h)<br>Group 2: flush with heparinized saline + WIT (160 ± 6.32 min) + SNMP with Steen+ (+19-21°C, 24h)<br>Control: flush with heparinized saline + SCS (4°C, 24h) | Group 1: Oxygenated Steen solution (7.5% albumin), albumin, glucose, ions vancomycin and piperacillin-tazobactam, sodium bicarbonate.<br>Group 2: Oxygenated Steen+ solution (15% albumin) albumin, glucose, ions vancomycin and piperacillin-tazobactam, sodium bicarbonate. Full-volume exchange after 12h. | Group 1: 6<br>Group 2: 6<br>Control: 6-12 | 18-24 | Group 1: more weight gain and edema (24.8% ± 5.5%); signs of compartment syndrome<br>Group 2: less weight gain (8.0 ± 4.7%); and stable dynamic/metabolic parameters; well-preserved tissues histologically and muscle contractility recovery |

CC3: Cleaved caspase 3; CIT: Cold ischemia time; CMAP: Compound muscle action potential; DDIT-4: DNA damage-inducible transcript 4; DMSO: Dimethyl sulfoxide; EVNLP: Ex vivo normothermic limb perfusion; H2S: Hydrogen sulfide solution; HBOC-201: Hemoglobin-based oxygen carrier-201; HIF1α: Hypoxia-inducible factor 1 alpha; HMGB-1: High mobility group box-1; HMP: Hypothermic Machine Perfusion; HSS: Histological severity score; HTK: Histidine–tryptophan–ketoglutarate solution; IT: Ischemia time; KPS-1: Kidney Preservation Solution-1 (UW); MO2: Dextran oxygen microcarriers; NMP: Normothermic Machine Perfusion; NO: Nitric oxide; NS: Normal saline; PEG: Polyethylene glycol / Polyethylene glycol-35; POD: Postoperative day; RBCs: Red blood cells; RGS-2: Regulator of G-protein signaling 2; RL: Ringer’s lactate solution; SCS: Static Cold Storage; SNMP: Su-bnormothermic Machine Perfusion; SSS: Supercooling storage solution; SZNF: Subzero nonfreezing; TIT: Total ischemia time; Txp: Transplantation; ULiSSES: Universal Limb Salvage System; UW: University of Wisconsin solution; WIT: Warm ischemia time.
Details regarding time, temperature, amount, or identities of solutions not listed were not reported by the study.

**Table 2.**
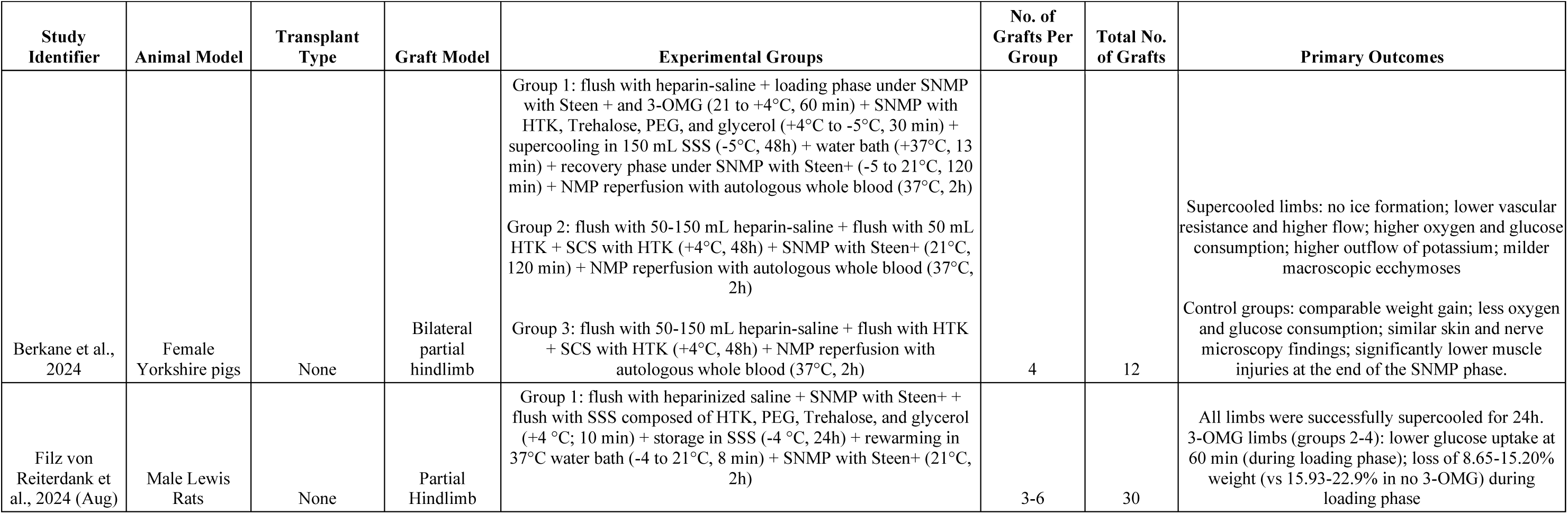
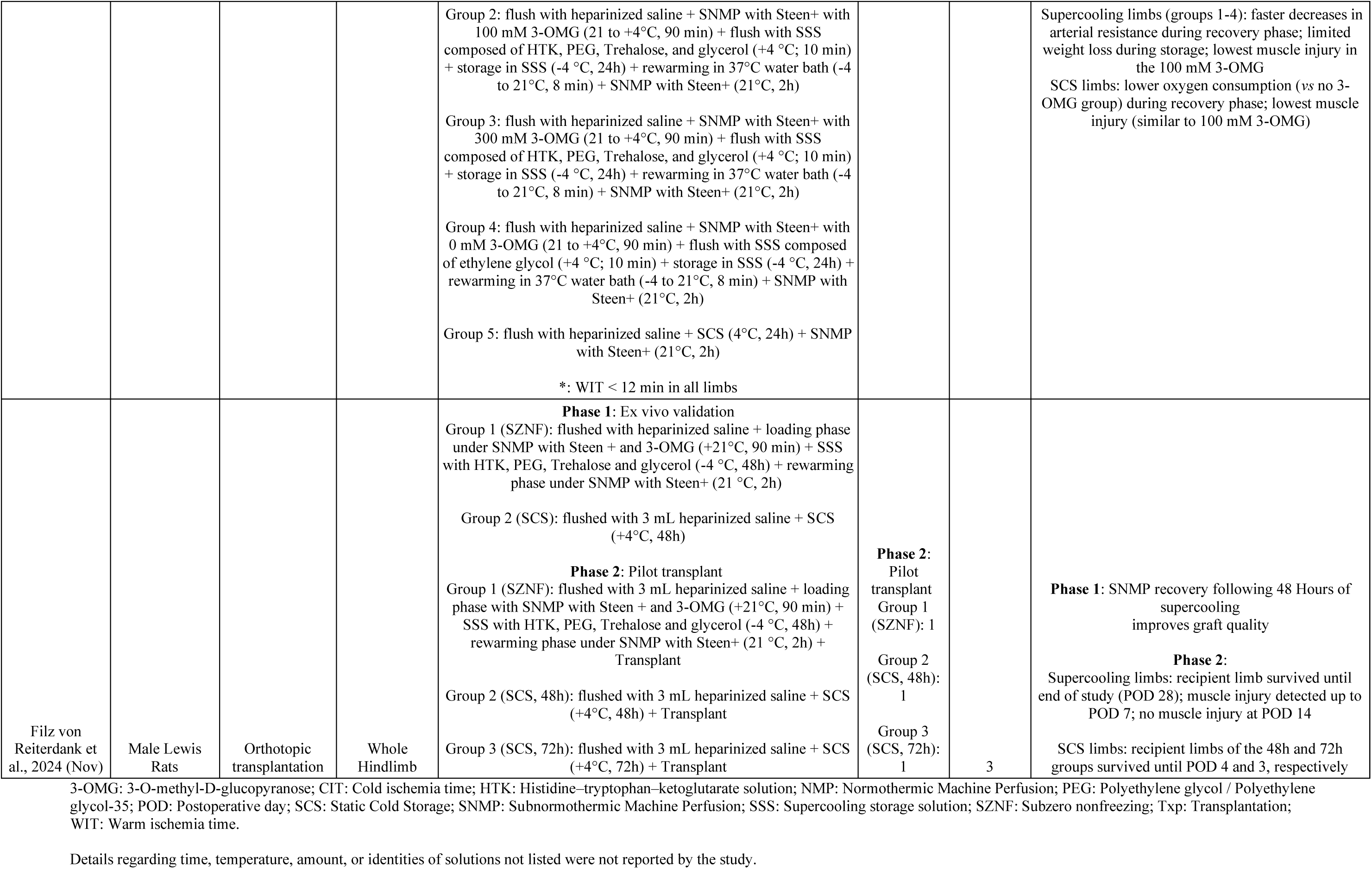
Characterizations of Cryopreservation Techniques.

| TABLE 2. Characterizations of Cryopreservation Techniques |  |  |  |  |  |  |  |
| --- | --- | --- | --- | --- | --- | --- | --- |
| Study Identifier | Animal Model | Transplant Type | Graft Model | Experimental Groups | No. of Grafts Per Group | Total No. of Grafts | Primary Outcomes |
| Berkane et al., 2024 | Female Yorkshire pigs | None | Bilateral partial hindlimb | Group 1: flush with heparin-saline + loading phase under SNMP with Steen + and 3-OMG (21 to +4°C, 60 min) + SNMP with HTK, Trehalose, PEG, and glycerol (+4°C to -5°C, 30 min) + supercooling in 150 mL SSS (-5°C, 48h) + water bath (+37°C, 13 min) + recovery phase under SNMP with Steen+ (-5 to 21°C, 120 min) + NMP reperfusion with autologous whole blood (37°C, 2h)<br>Group 2: flush with 50-150 mL heparin-saline + flush with 50 mL HTK + SCS with HTK (+4°C, 48h) + SNMP with Steen+ (21°C, 120 min) + NMP reperfusion with autologous whole blood (37°C, 2h)<br>Group 3: flush with 50-150 mL heparin-saline + flush with HTK + SCS with HTK (+4°C, 48h) + NMP reperfusion with autologous whole blood (37°C, 2h) | 4 | 12 | Supercooled limbs: no ice formation; lower vascular resistance and higher flow; higher oxygen and glucose consumption; higher outflow of potassium; milder macroscopic ecchymoses<br>Control groups: comparable weight gain; less oxygen and glucose consumption; similar skin and nerve microscopy findings; significantly lower muscle injuries at the end of the SNMP phase. |
| Filz von Reiterdank et al., 2024 (Aug) | Male Lewis Rats | None | Partial Hindlimb | Group 1: flush with heparinized saline + SNMP with Steen+ + flush with SSS composed of HTK, PEG, Trehalose, and glycerol (+4 °C; 10 min) + storage in SSS (-4 °C, 24h) + rewarming in 37°C water bath (-4 to 21°C, 8 min) + SNMP with Steen+ (21°C, 2h) | 3-6 | 30 | All limbs were successfully supercooled for 24h. 3-OMG limbs (groups 2-4): lower glucose uptake at 60 min (during loading phase); loss of 8.65-15.20% weight (vs 15.93-22.9% in no 3-OMG) during loading phase |
|  |  |  |  | Group 2: flush with heparinized saline + SNMP with Steen+ with 100 mM 3-OMG (21 to +4°C, 90 min) + flush with SSS composed of HTK, PEG, Trehalose, and glycerol (+4 °C; 10 min) + storage in SSS (-4 °C, 24h) + rewarming in 37°C water bath (-4 to 21°C, 8 min) + SNMP with Steen+ (21°C, 2h)<br>Group 3: flush with heparinized saline + SNMP with Steen+ with 300 mM 3-OMG (21 to +4°C, 90 min) + flush with SSS composed of HTK, PEG, Trehalose, and glycerol (+4 °C; 10 min) + storage in SSS (-4 °C, 24h) + rewarming in 37°C water bath (-4 to 21°C, 8 min) + SNMP with Steen+ (21°C, 2h)<br>Group 4: flush with heparinized saline + SNMP with Steen+ with 0 mM 3-OMG (21 to +4°C, 90 min) + flush with SSS composed of ethylene glycol (+4 °C; 10 min) + storage in SSS (-4 °C, 24h) + rewarming in 37°C water bath (-4 to 21°C, 8 min) + SNMP with Steen+ (21°C, 2h)<br>Group 5: flush with heparinized saline + SCS (4°C, 24h) + SNMP with Steen+ (21°C, 2h)<br>*: WIT < 12 min in all limbs |  |  | Supercooling limbs (groups 1-4): faster decreases in arterial resistance during recovery phase; limited weight loss during storage; lowest muscle injury in the 100 mM 3-OMG<br>SCS limbs: lower oxygen consumption (vs no 3-OMG group) during recovery phase; lowest muscle injury (similar to 100 mM 3-OMG) |
| Filz von Reiterdank et al., 2024 (Nov) | Male Lewis Rats | Orthotopic transplantation | Whole Hindlimb | **Phase 1:** Ex vivo validation<br>Group 1 (SZNF): flushed with heparinized saline + loading phase under SNMP with Steen + and 3-OMG (+21°C, 90 min) + SSS with HTK, PEG, Trehalose and glycerol (-4 °C, 48h) + rewarming phase under SNMP with Steen+ (21 °C, 2h)<br>Group 2 (SCS): flushed with 3 mL heparinized saline + SCS (+4°C, 48h)<br>**Phase 2:** Pilot transplant<br>Group 1 (SZNF): flushed with 3 mL heparinized saline + loading phase with SNMP with Steen + and 3-OMG (+21°C, 90 min) + SSS with HTK, PEG, Trehalose and glycerol (-4 °C, 48h) + rewarming phase under SNMP with Steen+ (21 °C, 2h) + Transplant<br>Group 2 (SCS, 48h): flushed with 3 mL heparinized saline + SCS (+4°C, 48h) + Transplant<br>Group 3 (SCS, 72h): flushed with 3 mL heparinized saline + SCS (+4°C, 72h) + Transplant | **Phase 2:** Pilot transplant<br>Group 1 (SZNF): 1<br>Group 2 (SCS, 48h): 1<br>Group 3 (SCS, 72h): 1 | 3 | **Phase 1:** SNMP recovery following 48 Hours of supercooling improves graft quality<br>**Phase 2:**<br>Supercooling limbs: recipient limb survived until end of study (POD 28); muscle injury detected up to POD 7; no muscle injury at POD 14<br>SCS limbs: recipient limbs of the 48h and 72h groups survived until POD 4 and 3, respectively |

3-OMG: 3-O-methyl-D-glucopyranose; CIT: Cold ischemia time; HTK: Histidine–tryptophan–ketoglutarate solution; NMP: Normothermic Machine Perfusion; PEG: Polyethylene glycol / Polyethylene glycol-35; POD: Postoperative day; SCS: Static Cold Storage; SNMP: Subnormothermic Machine Perfusion; SSS: Supercooling storage solution; SZNF: Subzero nonfreezing; Txp: Transplantation; WIT: Warm ischemia time.
Details regarding time, temperature, amount, or identities of solutions not listed were not reported by the study.

### Static Cold Storage

Only one study examined static cold storage (SCS) as the main preservation technique. In their experiment on transplanted rat hindlimb, Goutard et al. evaluated the possibility of extending the duration of SCS by using sub-normothermic machine perfusion (SNMP) as a recovery adjunct to SCS. After determining that the optimal SCS duration in a rodent hindlimb transplant model was 12h, the authors compared limbs transplanted directly after 12h of SCS with limbs transplanted following 3h of recovery with SNMP. The SNMP limbs showed faster recovery from ischemic lesions and no or minimal degeneration on biopsies performed on postoperative day (POD) 21. However, no statistical difference in graft survival was observed between the two groups. The authors concluded that this hybrid method can prolong the viability of the limb stored under SCS past the 6h mark (17).

### Ex vivo Machine Perfusion

Broadly speaking, machine perfusion (MP) can be classified, based on perfusion temperatures, into hypothermic (0 °C – 12 °C), mid-thermic MP (13°C–24°C); sub-normothermic (25 °C – 34 °C), and normothermic or physiologic (35°C–38°C) (31). In the articles reviewed, 4 studies used normothermic MP (NMP) (15,18,22,24), 5 studies used sub-normothermic MP (SNMP) (14,19–21,27), and 4 studies used hypothermic MP (HMP) (16,23,25,26).

#### Normothermic Machine Perfusion (NMP)

NMP maintains normal cellular metabolism at physiological temperatures thereby reducing ischemia-reperfusion injury (IRI) (32). In a canine model, Chen et al. compared 24h of NMP with an oxygenated autologous blood perfusate to SCS. Although NMP limbs gained more weight than SCS limbs (11.72% ± 4.06% *vs* 0.18% ± 0.10%) and histology showed superior preservation of muscle architecture with no significant change from baseline in the NMP group, leading the authors to conclude that NMP with autologous blood can sustain limb viability for at least 24 h (24). In the only human study, 10 human forearms perfused with oxygenated autologous blood under NMP were compared with their contralateral counterparts preserved with SCS. After an average NMP duration of 41.6 ± 9.4h, the authors found that NMP with autologous blood allows extended preservation times while preserving muscle contractility until a median of 30.5h (15). Despite these promising results, red blood cell (RBC)-based perfusates have limitations, including limited availability, cross-matching requirements, and potential pro-inflammatory effects (18,33). To address these issues, alternative oxygen carriers have been investigated (18,22,33). Figueroa et al. found that a polymerized hemoglobin-based oxygen carrier-201 (HBOC-201) perfusate performed comparably to RBC-based perfusates in supporting porcine limb physiology and metabolism (18). Likewise, Haug et al. demonstrated that acellular dextran oxygen microcarriers in NMP of rat hindlimbs maintained mitochondrial structure and reduced muscle injury markers compared with conventional acellular perfusates (22).

#### Sub-Normothermic Machine Perfusion (SNMP)

While lowering the temperature by 10°C can reduce metabolism by as much as 50%, cold-induced injury remains a risk (1,24,34). Thus, SNMP offers an intermediate metabolic state between NMP and HMP. In 2024, Goutard et al. compared 24 h SNMP to SCS in porcine partial hindlimb allotransplantation: two of three SCS recipients died before POD 14, while all SNMP recipients survived with viable grafts, confirming SNMP’s ability to maintain tissue viability (21). Perfusate composition is another critical factor. Oubari et al. showed that Steen+ (15% albumin) reduced edema (8.0% ± 4.7% *vs* 24.8% ± 5.5% of mean weight gain) and histologic injury compared with standard Steen (7.5%) while allowing muscular contraction recovery (2.4/5 ± 0.89 *vs* 0/5 after 24 h) (14). Flow and pressure parameters have also been explored. A porcine study found continuous flow provided more stable resistances, while pulsatile flow enhanced vasoconductivity and reduced ischemic injury (20). Veraza et al. tested a closed vertical SNMP device (ULiSSES), which reduced weight gain by 21%, decreased intracellular edema, and increased oxygen consumption compared with an open system, while offering portability for remote use (19). Finally, Charlès et al. demonstrated that nonrecirculating SNMP can restore viability after up to 120 min of warm ischemia in rat hindlimbs. Post-replantation, 80% of controls developed necrosis, whereas no necrosis occurred in the SNMP group, an effect attributed to toxin clearance, improved oxygen use, and reduced lactate (27).

#### Hypothermic Machine Perfusion (HMP)

An even more drastic reduction in metabolic demand can be expected with HMP offering the potential to further extend the preservation window in VCA. In single-cohort study, Zhang et al. successfully replanted six porcine forelimbs after 9 hours of warm ischemia at 22°C followed by 24 hours of HMP at 8 to 10 °C using an oxygenated modified histidine-tryptophanketoglutarate (HTK) solution. Only mild histological signs of muscle injury were observed 12 h post-replantation, leading the authors to conclude that HMP can safely extend preservation up to 33 h (26). Similarly, Brouwers et al. noted minimal degeneration of muscle (histologic injury severity score [HISS] of 1.8 in SCS and 3.8 in HMP) seven days post-replantation of porcine forelimbs preserved under HMP with oxygenated HTK for 24h. Although HMP limbs showed increased edema (2% decrease in weight in SCS *vs* 44% increase in HMP) and variation in muscle cells compared with the limbs preserved with 4h of SCS, overall graft viability remained comparable between groups (25). Perfusate composition has also been examined under hypothermic conditions. In 2022, Brouwers et al, using 4h SCS as a control, compared HTK with UW for the 24h preservation at 8°C of porcine abdominal myocutaneous flaps before replantation. While the outcomes were macroscopically comparable on POD 7, flaps perfused with HTK demonstrated less histologic muscle damage compared to UW-perfused flaps (16). More recently, Ng et al compared HMP with University of Wisconsin Kidney Preservation Solution-1 (KPS-1) versus normal saline (NS) in a rat hindlimb model. After 24h at 4°C, KPS-1 limbs had a significantly lower weight gain than the NS-preserved limbs (13.3% ± 0.04% *vs* 69.5% ± 0.17%; P < 0.001) making them suitable candidates for orthotopic transplantation whereas the NS limbs were not due to excessive edema (23).

### Supercooling

Since its first description by Nakagawa et al in 1997 (35), subzero non-freezing preservation has gained considerable attention in the transplantation sciences (30,36–38). The technique aims to decrease cell metabolism through lower temperatures while avoiding ice nucleation (30). This review identified three articles applying supercooling in VCA preservation to rodent (28,29) and swine models (39). In 2024, Filz von Reiterdank et al. introduced a supercooling protocol using SNMP for both loading and recovery. In rodent partial hindlimbs preserved for 24 h, supercooled grafts demonstrated faster decreases in arterial resistance (mean of 272.7 ± 132.1 mmHg(min/mL) at 60 min in supercooling *vs* 717.5 ± 164.2 mmHg(min/mL) in SCS; P < 0.001) and a trend toward weight loss, whereas SCS limbs gained weight. No significant histologic differences were observed, but the findings suggested that supercooling could extend VCA preservation (29). The same group later evaluated whole rodent hindlimbs supercooled at –4 °C for 48 h (28). After 48h of supercooling, the hindlimbs showed improved oxygen uptake, glucose consumption, and lower vascular resistance, with skin architecture preserved and only moderate muscle ischemia confirming graft recovery. Despite weight gain of the supercooled limb exceeding 30%, considered a limit for VCA transplantability in partial hindlimb model, the authors performed a pilot study comparing orthotopic transplantation of a hindlimb preserved with 24h supercooling versus SCS for either 48h or 72h. The recipient of the supercooled limb survived until the end of the study (POD 28) while the recipients of the limbs preserved with SCS for 48h and 72h only survived until POD 4 and 3, respectively. At the end of the study, the supercooled graft showed patent vessels, unremarkable skin and muscle biopsies, and a return to its normal size (28). Building on this work, Berkane et al. tested supercooling in a large-animal model. Four porcine partial hindlimbs were stored at –5 °C for 48 h, with CPA loading via 1h SNMP pre-supercooling and 2 h SNMP recovery, followed by 2 h NMP. Compared with the two control groups preserved by SCS with or without SNMP recovery, supercooled grafts showed improved vascular resistance, higher oxygen consumption, and less edema, with better muscle preservation on histology. Importantly, no ice formation was observed, demonstrating the feasibility of 48 h supercooling in large-animal VCAs (39).

### Risk of Bias assessment

Quality assessment revealed an overall low methodological level across the 16 preclinical animal studies (14,16–29,39) evaluated using SYRCLE (12), reflecting the high risk of selection and performance bias. These shortcomings were largely attributable to the lack of allocation concealment and absence of randomization in group assignment. Most studies demonstrated low risk in other bias domains, with a few exceptions: Veraza et al., where a potential conflict of interest existed as the authors work for the device manufacturer; Chen et al. and Ng et al., where assessor blinding was insufficiently reported; and Brouwers et al. (2024), where handling of missing data was unclear. In contrast, the single human ex vivo study (15) demonstrated a low risk of bias scoring “Yes” across all the items of the JBI appraisal checklist (13), underscoring its methodological robustness. Details on the risk of bias assessment of included studies are reported in Supplemental material 2.

## Discussion

This systematic review identified 17 studies on ex vivo VCA preservation, compared with 13 in our previous review, reflecting the rapid growth of the field in the past three years. Preservation strategies were grouped into static cold storage (SCS), ex vivo machine perfusion (MP), and supercooling. Since the last review, only one new SCS study (17) has been published, while 13 addressed on machine perfusion (14–16,18–27) and 3 investigated supercooling (28,29,39). The decline in SCS-related work suggests waning interest in this approach and a growing focus on machine perfusion, consistent with trends in solid organ transplantation (40–42). The SCS study evaluated of SNMP as recover method after extended cold storage (17), and supercooling protocols incorporated SNMP during both loading and recovery phases (28,29,39). Whereas earlier studies focused on establishing the feasibility of ex vivo perfusion in VCA (10), recent research has shifted towards optimization testing parameters such as temperature, perfusate composition, flow, and pressure (14–16,18–27). Although not a novel approach, supercooling is attracting increasing interest in the field of VCA preservation.

### Static Cold Storage

Cold flush followed by static cold storage (SCS) at 4°C remains the gold standard for VCA preservation. Earlier research in this field primarily sought to optimize storage solutions in order to mitigate the ischemic injury associated with SCS (10,43,44). While new cold-storage solutions have been developed and applied in solid organ transplantation (45), no recent studies have specifically investigated alterations to SCS solutions in VCA. Instead, Goutard et al. (17) employed a hybrid approach, combining 12h of SCS with an oxygenated modified-Steen solution followed by 3h SNMP for rodent hindlimb recovery. This approach allowed controlled rewarming, clearance of accumulated toxic metabolites, and normalization of lactate and potassium levels to those seen in a fresh grafts. While graft survival did not significantly differ, SNMP-treated limbs exhibited more rapid recovery from ischemic injury and reduced histological damage in muscle and skin (17). These findings suggest that applying SNMP after SCS could be a promising method to recondition grafts and extend preservation beyond the conventional 6h threshold. This concept aligns with evidence from solid organ transplantation, where machine perfusion has successfully prolonged SCS duration and improved post-transplant outcomes. (46,47).

### Ex vivo Machine Perfusion

First introduced in the 1930s (48,49) and applied clinically to liver and kidney transplantation in the 1960s (50,51), machine perfusion (MP) initially fell out of favor before experiencing a resurgence with the first modern clinical series in 2010 (52). Since then, MP has gained widespread adoption, culminating in FDA approval of the first ex vivo liver perfusion device in 2021 (53). With its safety and efficacy well established in solid organ transplantation, VCA research now focuses on determining the best approach to translate MP into clinical practice (54).

Perfusion temperature and solution composition remain the two principal variables distinguishing MP systems. Among the included studies, SNMP was the most common approach (14,19–21,27), followed by NMP (15,18,22,24) and HMP (16,23,25,26). No study directly compared perfusion temperatures. A wide range of perfusates was used: blood-based perfusates (15,24); colloid-based solutions like UW (16) and Steen (14,17,20–22,27); and crystalloid-based solutions like HTK (16,25,26), KPS-1 (23), Plasma-Lyte (18), Krebs–Henseleit (19), and normal saline (23). Oxygenation is required in NMP and seems to carry an advantage in perfusion at lower temperatures (55,56). However, reliance on RBC-based perfusates is limited by issues of availability, infection risk, and immunogenicity. To overcome these challenges, alternative oxygen carriers have been explored (57,58). Figueroa et al. demonstrated that a polymerized hemoglobin-based oxygen carrier-201 (HBOC-201) could maintain metabolism and function in porcine limbs comparably to RBC-based perfusates (18), while Haug et al. reported that dextran-based oxygen microcarriers supported mitochondrial integrity and reduced muscle injury markers relative to conventional acellular perfusates (22). Nevertheless, both studies observed substantial weight gain, with increases up to 124% in limbs perfused with Phoxilium enriched with oxygen carriers (22). These findings highlight the promise of hemoglobin-based and synthetic carriers as substitutes for RBCs, but also the need for further optimization. Another common strategy to optimise perfusate composition is periodic replacement of the solution to avoid accumulation of toxic metabolites (14,15,18,20,21,24). Charlès et al. recently showed that a nonrecirculating oxygenated modified Steen solution can restore rat hindlimb viability after prolonged warm ischemia (27).

Two other important MP parameters that vary between species and graft type are pressure and flow (54,59). Perfusion pressures varied across studies, and no study has directly compared different MP pressures in VCA preservation. One study compared a pressurized perfusion system (ULiSSES) to the standard nonpressurized MP and found a decrease in weight gain and edema, along with an increase in oxygen consumption and flow after 24h of SNMP (19). This design also offered greater portability, potentially facilitating graft recovery in remote settings. Regarding flow dynamics, some evidence suggests pulsatile flow (PF) may enhance reconditioning (60) and attenuate inflammatory responses compared with continuous flow (CF) (61) but no consensus has been found (20,56). In a swine hindlimb model, Tawa et al. compared PF (at 60 beat per minute) with CF during 24-h SNMP and observed improved endothelial function and reduced ischemic injury (20). Further studies isolating the effects of pressure and flow are needed to identify optimal configurations.

Despite these advances, challenges remain. Graft weight gain, a consistent finding across multiple studies (16,18–24,26), represents a key barrier to clinical translation. Weight gain is recognised as an early marker of graft injury and may preclude transplantation (62). Strategies to address this issue include increasing oncotic pressure, as shown by Oubari et al. with albumin-enriched Steen solution (14) and modifying perfusion mechanics, as in Veraza’s pressurized system (19). Nonetheless, the ideal combination of temperature, perfusate, and circuit design has yet to be defined. Even so, the body of evidence demonstrates that ex vivo perfusion remains a highly promising avenue toward overcoming the preservation barriers that currently limit the clinical expansion of VCA.

### Supercooling

In addition to SCS, other static preservation strategies have been explored. Our previous review (10) identified one study on cryopreservation, in which graft survival was maintained following storage at ultralow temperatures and subsequent thawing (63). Since then, however, no further studies have investigated cryopreservation in the context of VCA. More recently, attention has shifted toward supercooling, or subzero nonfreezing preservation, which offers the potential for multi-day storage and could facilitate the implementation of immune tolerance protocols in recipients (30). Supercooling requires coupling with SNMP, both as a loading phase to introduce protective agents and as a recovery phase after storage (3). Initially studied in the liver, where preservation up to 96 h was achieved in rat models (47), supercooling has only recently been applied to VCA. To date, three studies from the same research team have reported encouraging results in rodent and porcine models (28–30). Most notably, in a pilot study on whole rat hindlimbs, Filz von Reiterdank et al. successfully transplanted a limb preserved for 48h at –4 °C, with survival observed up to 28 days postoperatively (28). While further optimization - particularly of storage solutions - is still required, preliminary data suggest that supercooling may play a crucial role in prolonging preservation times and improving outcomes in VCA.

### Limitations

This review has several limitations, many of which stem from the current state of the literature on VCA preservation. Although the number of studies on preservation strategies has increased in recent years, the available primary evidence remains limited. Consequently, heterogeneity in methods and outcome reporting precludes reliable statistical analysis. All included studies were preclinical, relying mainly on animal models with small sample sizes and short follow-up durations, which restricts extrapolation to the clinical setting. Moreover, only a minority of studies assessed preserved VCAs in vivo through transplantation or replantation, making it difficult to fully appreciate the physiological impact of the interventions. Functional outcomes themselves were rarely evaluated, despite being critical for long-term graft success beyond simple viability. Another limitation is the lack of consideration for donor types which do not reflect clinical transplantation, where donor physiology critically influences ischemia tolerance and preservation outcomes. VCA models were almost exclusively procured from living anesthetized animals, without simulating brain-death or circulatory-death donors (Maastricht classification), or considering donor criteria such as age and comorbidities. Finally, while our primary objective was to evaluate cellular physiology, it must be acknowledged that immunologic factors play a pivotal role in VCA success and warrant further study. Future research should employ standardized protocols and prioritize long-term functional and immunologic outcomes to better assess the effectiveness of different preservation strategies in transplanted VCAs.

## Conclusion

This review highlights the rapid growth of research on VCA preservation methods. While SCS remains the clinical standard, recent studies emphasize machine perfusion techniques and hybrid strategies that optimize perfusate composition, pressure/flow dynamics, and circuit design to extend preservation and reduce ischemic injury. Supercooling, combined with SNMP loading and recovery, is being explored and has shown promising results for multi-day storage in both rodent and porcine models. Despite these advances, most studies remain preclinical with small sample sizes, methodological differences, and limited functional or immunologic endpoints, making translation to clinical practice premature. Future research should standardize protocols, incorporate transplantation outcomes, and focus on large-animal and early human studies to establish preservation strategies that reliably extend graft viability and function.

## Supporting information

Supplementary Material 1. Database Search Strategies.pdf

Supplemental material 2. Risk of Bias Evaluation for Included Studies

## Author Contributions

Conceptualization, E.L.; methodology, E.L. P.B.P.N.; software, P.N.; validation, P.B. and E.L.; formal analysis, P.B. and P.N.; investigation, P.N.; resources, E.L, O.C, E.R.; data curation, P.N.; writing—original draft preparation, P.N.; writing—review and editing, E.L., O.C., P.N, P.B., E.R, D.P., P.R., A.S. visualization, E.R, E.L.; supervision, P.R, E.L., E.R.; project administration, E.L.; funding acquisition, E.L. All authors have read and agreed to the published version of the manuscript.

## Institutional Review Board Statement

This systematic review was prospectively approved and was registered by the Institutional Review Board of the University Institute for Locomotion and Sport with the No. IRB00014528_2025_23.

## Informed Consent Statement

Patient consent was not required as this study is a systematic review of previously published literature.

## Data Availability Statement

The corresponding authors can provide the data upon request.

**Supplementary material 1. Database Search Strategies**

**Supplementary material 2. Risk of Bias Evaluation for Included Studies**

