## Supplementary Material 1. Database Search Strategies.pdf for "Preservation Strategies for Vascularized Composite Allotransplantation: An Updated Systematic Review of a Rapidly Expanding Field"

**Table 1. Embase search**

| No. | Query | Results |
|---|---|---|
| #6 | #5 AND (2022:py OR 2023:py OR 2024:py OR 2025:py) | 916 |
| #5 | 'face transplant'/exp OR 'face transplant' OR 'facial transplant'/exp OR 'facial transplant' OR 'face transplantation'/exp OR 'face transplantation' OR 'facial transplantation'/exp OR 'facial transplantation' OR 'face allotransplantation' OR 'facial allotransplantation' OR 'facial vascularized composite allotransplantation' OR 'face vascularized composite allotransplantation' OR 'face vascularized composite allograft' OR 'facial vascularized composite allograft' OR 'face allograft' OR 'facial allograft' OR 'face composite tissue allotransplantation' OR 'facial composite tissue allotransplantation' OR 'face composite tissue allograft' OR 'facial composite tissue allograft' OR 'hand allograft' OR 'hand composite tissue allotransplantation' OR 'hand composite tissue allograft' OR 'hand transplantation'/exp OR 'hand transplantation' OR 'hand transplant' OR 'hand allotransplantation' OR 'hand vascularized composite allotransplantation' OR 'hand vascularized composite allograft' OR 'vascularized allotransplantation' OR 'vascularized composite allotransplantation'/exp OR 'vascularized composite allotransplantation' OR 'vascularized composite allograft'/exp OR 'vascularized composite allograft' OR 'vascularized allograft' OR 'composite tissue allotransplantation'/exp OR 'composite tissue allotransplantation' OR 'composite tissue allograft'/exp OR 'composite tissue allograft' OR 'vascularized tissue allotransplantation' OR 'vascularized tissue allograft' | |

**Table 2. PuMed/Medline search**

("facial transplantation"[Title/Abstract] OR "face transplant"[Title/Abstract] OR "facial transplant"[Title/Abstract] OR "face transplantation"[Title/Abstract] OR "facial transplantation"[Title/Abstract] OR "face allotransplantation"[Title/Abstract] OR "facial allotransplantation"[Title/Abstract] OR "facial vascularized composite allotransplantation"[Title/Abstract] OR "face vascularized composite allotransplantation"[Title/Abstract] OR "face vascularized composite allograft"[Title/Abstract] OR "facial vascularized composite allograft"[Title/Abstract] OR "face allograft"[Title/Abstract] OR "facial allograft"[Title/Abstract] OR "face composite tissue allotransplantation"[Title/Abstract] OR "facial composite tissue allotransplantation"[Title/Abstract] OR "face composite tissue allograft"[Title/Abstract] OR "facial composite tissue allograft"[Title/Abstract] OR "hand allograft"[Title/Abstract] OR "hand composite tissue allotransplantation"[Title/Abstract] OR "hand composite tissue allograft"[Title/Abstract] OR "hand transplantation"[Title/Abstract] OR "hand transplant"[Title/Abstract] OR "hand allotransplantation"[Title/Abstract] OR "hand vascularized composite allotransplantation"[Title/Abstract] OR "hand vascularized composite allograft"[Title/Abstract] OR "vascularized allotransplantation"[Title/Abstract] OR "vascularized composite allotransplantation"[Title/Abstract] OR "vascularized composite allograft"[Title/Abstract] OR "vascularized allograft"[Title/Abstract] OR "composite tissue allotransplantation"[Title/Abstract] OR "composite tissue allograft"[Title/Abstract] OR "vascularized tissue allotransplantation"[Title/Abstract] OR "vascularized tissue allograft"[Title/Abstract])

**Table 3. Cochrane search**

1 Trial matching ("facial transplantation" OR "face transplant" OR "facial transplant" OR "face transplantation" OR "facial transplantation" OR "face allotransplantation" OR "facial allotransplantation" OR "facial vascularized composite allotransplantation" OR "face vascularized composite allotransplantation" OR "face vascularized composite allograft" OR "facial vascularized composite allograft" OR "face allograft" OR "facial allograft" OR "face composite tissue allotransplantation" OR "facial composite tissue allotransplantation" OR "face composite tissue allograft" OR "facial composite tissue allograft" OR "hand allograft" OR "hand composite tissue allotransplantation" OR "hand composite tissue allograft" OR "hand transplantation" OR "hand transplant" OR "hand allotransplantation" OR "hand vascularized composite allotransplantation" OR "hand vascularized composite allograft" OR "vascularized allotransplantation" OR "vascularized composite allotransplantation" OR "vascularized composite allograft" OR "vascularized allograft" OR "composite tissue allotransplantation" OR "composite tissue allograft" OR "vascularized tissue allotransplantation" OR "vascularized tissue allograft")
